# Effects of dopamine and opioid receptor antagonism on the neural processing of social and non-social rewards

**DOI:** 10.1101/2023.06.16.545306

**Authors:** Claudia Massaccesi, Sebastian Korb, Sebastian Götzendorfer, Emilio Chiappini, Matthaeus Willeit, Johan N. Lundström, Christian Windischberger, Christoph Eisenegger, Giorgia Silani

## Abstract

Rewards are a broad category of stimuli inducing approach behavior to aid survival. Extensive evidence from animal research has shown that wanting (the motivation to pursue a reward) and liking (the pleasure associated with its consumption) are mostly regulated by dopaminergic and opioidergic activity in dedicated brain areas. However, less is known about the neuroanatomy of dopaminergic and opioidergic regulation of reward processing in humans, especially when considering different types of rewards (i.e., social and non-social). To fill this gap of knowledge, we combined dopaminergic and opioidergic antagonism (via amisulpride and naltrexone administration) with functional neuroimaging to investigate the neurochemical and neuroanatomical bases of wanting and liking of matched non-social (food) and social (interpersonal touch) rewards, using a randomized, between-subject, placebo-controlled, double-blind design. While at the behavioral level no drug effect was observed, brain activity was modulated by the administered compounds. In particular, opioid antagonism, compared to placebo, was associated with reduced activity in the medial orbitofrontal cortex during consumption of the most valued social and non-social rewards. Dopamine antagonism, however, had no clear effects on brain activity in response to rewards anticipation. These findings provide insights into the neurobiology of human reward processing and suggest a similar opioidergic regulation of the neural responses to social and non-social reward consumption.

## Introduction

Rewards are powerful drivers of human behavior. They guide our motivation and choices, on which our wellbeing and survival depend. Research has revealed at least two main dissociable reward-related components: *wanting* (or incentive salience), i.e., the motivation to pursue a reward, and *liking*, i.e., the hedonic aspect associated with its consumption (Berridge & Kringelbach, 2015; Berridge & Robinson, 2003). Thirty years of animal research (Berridge & Kringelbach, 2015; Berridge & Valenstein, 1991; Peciña et al., 2006; Peciña & Smith, 2010), as well as recent behavioral findings in humans (Buchel et al., 2018; Chelnokova et al., 2014; Eikemo et al., 2016; Korb, Götzendorfer, et al., 2020; but see also Soutschek et al., 2021), indicate that the wanting-liking dissociation is reflected at the neuroanatomical and neurochemical level. Dopaminergic activity in the midbrain and striatum directly subserves wanting during reward anticipation. Opioidergic activity in the nucleus accumbens (NAc) regulates specifically the hedonic reactions (liking) linked to reward consumption, and, indirectly, reward motivation. While such dissociation has been extensively documented in anticipation and consumption of palatable food and addictive drugs in animal studies, evidence for other types of rewards is scarce, especially in humans, and it remains unclear whether social and non-social rewards rely on the same neuroanatomical and neurochemical bases.

Previous human neuroimaging studies provided evidence for the presence of an “extended common currency schema” (Ruff & Fehr, 2014) in the brain (Gu et al., 2019; Izuma et al., 2008; Lin et al., 2012; Liu et al., 2011; Sescousse et al., 2013; Wake & Izuma, 2017). According to this model, reward magnitude is processed in the same reward-related brain structures, such as the striatum or the ventromedial prefrontal cortex (vmPFC), independently of the type of reward. However, differences in the neural responses to social and non-social rewards have also been documented, suggesting, for instance, reward-specific representations in the orbitofrontal cortex (OFC) (Sescousse et al., 2010). These findings point to the existence of a dedicated neural circuitry for the encoding of social reward values (“social valuation specific schema”; Ruff & Fehr, 2014), which may be selectively impaired in conditions characterized by deficits in social reward processing, like autism (Chevallier et al., 2012). Further, it was recently proposed that while domain-general neural circuits underlie reward anticipation (Gu et al., 2019), modality-specific ones may be involved during reward consumption (Rademacher et al., 2010). The incongruity of the reported results may stem from various methodological pitfalls, such as assessing exclusively reward anticipation but not consumption or employing social and non-social stimuli which are not matched in terms of primacy and magnitude, hence not accounting for possible differences in the properties of the rewards. Notably, by using a paradigm tailored to investigate anticipation and consumption of well-matched social and non-social primary rewards (Chiappini et al., 2022), we recently observed partially different effects of dopaminergic and opioidergic pharmacological challenges during anticipation of food and social touch rewards (Korb, Götzendorfer, et al., 2020). Consistent with animal studies, we showed that opioid antagonism modulates both anticipation and consumption of rewards, while dopaminergic antagonism affects only reward anticipation. However, the reduction in wanting following dopamine and opioid antagonism was stronger or limited to the anticipation of food rewards. Crucially, how these neurochemical mechanisms are reflected at the neurophysiological and neuroanatomical levels is still an open question.

By combining pharmacology with functional neuroimaging, we aimed to fill this knowledge gap and provide novel insights on the human neurochemical and neuroanatomical bases of the motivational and hedonic aspects of social and non-social reward processing. Specifically, in a randomized, double-blind, placebo-controlled, between-subjects design, 89 participants received orally either the highly selective D2/D3 dopamine receptor antagonist amisulpride (400 mg), the non-selective opioid receptor antagonist naltrexone (50 mg), or a placebo pill (see Table 1 for details about the sample’s characteristics). Drug effects on behavioral and neural responses to anticipation and consumption of food (milk with three different concentrations of chocolate) and interpersonal non-sexual touch (caresses to the forearm at three different speeds by a same-gender confederate) were assessed on a trial-by-trial basis, using implicit (i.e., physical effort) and explicit (subjective ratings of wanting and liking) measures (Chiappini et al., 2022; see Figure 1). Rewarding stimuli were carefully matched in terms of magnitude (similar levels of wanting and liking as shown in previous studies; Korb, Götzendorfer, et al., 2020; Korb, Massaccesi, et al., 2020), primacy (both food and touch can be classified as primary rewards, in contrast to money or social feedback), temporal proximity, tangibility, and familiarity (Matyjek et al., 2020).

**Figure 1.**
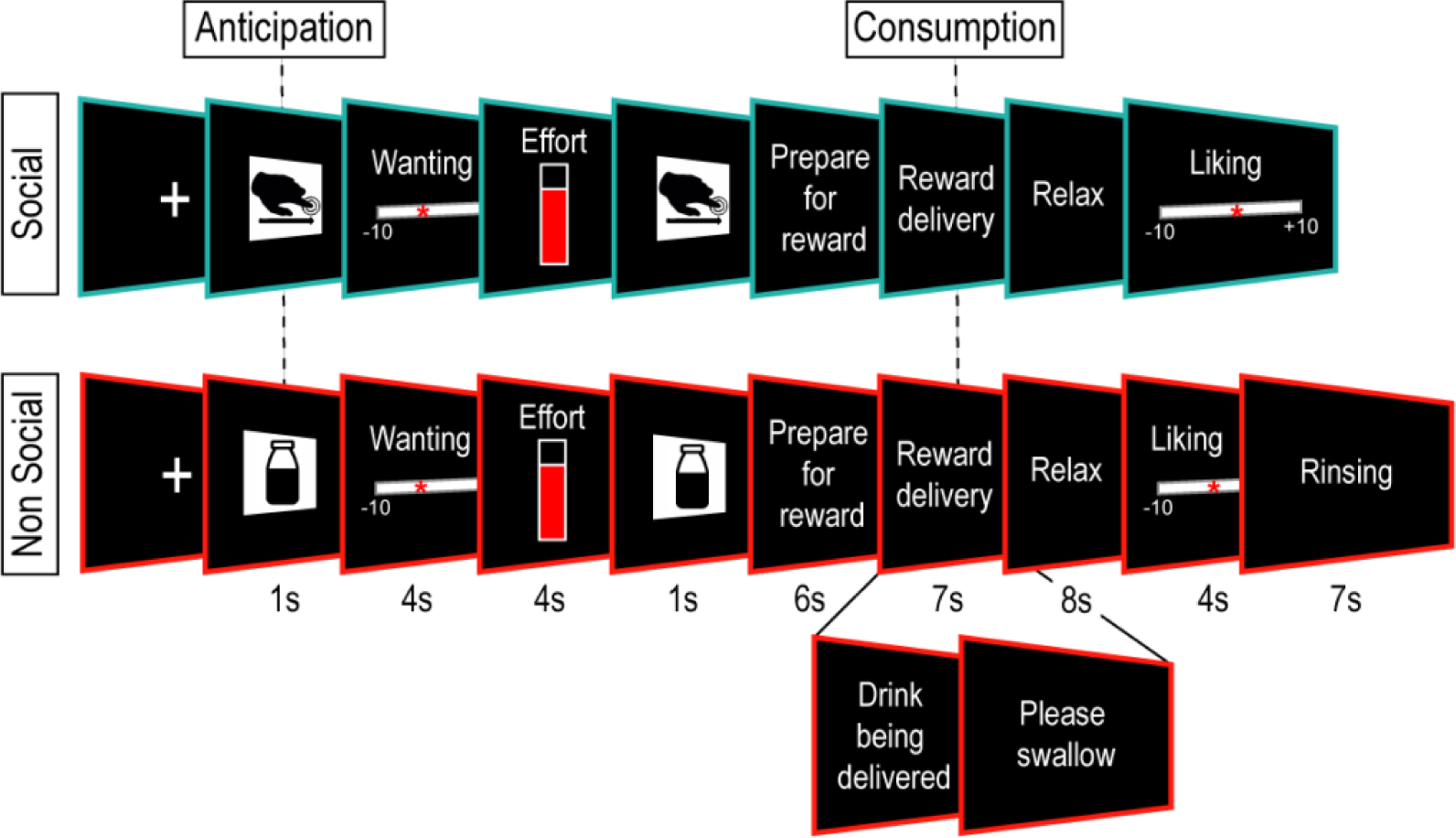
Trial sequence of the reward task. Participants could obtain social and non-social rewards in three levels (high, low, very low). Social rewards consisted of skin-to-skin caresses delivered to the forearm from a trained same-gender confederate at three speeds: 6 cm/s, 21 cm/s, and 27 cm/s. Non-social rewards consisted of milk with three different concentrations of cocoa: chocolate milk, a 4:1 mix of milk and chocolate milk, and milk. At the beginning of each trial, a cue announced the attainable reward (high or low; 1s), and participants were asked to rate their wanting of the announced reward (4s). Then participants exerted effort by squeezing a hand dynamometer to obtain the announced reward (4s). The applied force, displayed via online visual feedback, was expressed as percentage of the participants’ maximum voluntary contraction (MVC, measured immediately before the task) and translated into the probability of obtaining the announced reward (0 – 100%). The obtained reward was then announced (high, low, or, in case of low effort, very low; 1s) and delivered (7s; in non-social trials, participants were instructed to swallow following 2.5 s of drink delivery). Following a relaxation phase (8s), participants rated their liking of the stimulus (4s). At the end of non-social trials, participants received water to rinse their mouth (7s).

**Table 1.** Sample characteristics.

|  | Amisulpride | Naltrexone | Placebo | Group differences |  |
| --- | --- | --- | --- | --- | --- |
|  |  |  |  | *F* | *p* |
| N (Male, Female) | 32 (14, 18) | 29 (10, 19) | 28 (11, 17) | - | - |
| Age | 24.0 ± 4.0 | 23.9 ± 3.6 | 23.3 ± 3.6 | .36 | .70 |
| BMI | 23.0 ± 2.6 | 22.3 ± 2.2 | 22.8 ± 2.2 | .83 | .44 |
| MVC pre | 153.5 ± 56.8 | 153.6 ± 62.8 | 139.9 ± 54.8 | .50 | .61 |
| MVC post | 169.2 ± 54.2 | 160.5 ± 58.0 | 160.8 ± 53.5 | .23 | .80 |
| Hunger | 3.2 ± 2.0 | 3.6 ± 2.0 | 3.5 ± 4.8 | .36 | .70 |
| Sociability | 5.0 ± 1.2 | 4.8 ± 1.7 | 4.8 ± 1.3 | .13 | .88 |
| AQ-k | 7.2 ± 4.7 | 5.4 ± 3.1 | 5.5 ± 3.1 | 2.47 | .09 |
| HTAS | 66.3 ± 14.7 | 72.4 ± 13.7 | 67.4 ± 13.7 | 1.82 | .17 |
| STQ | 22.7 ± 8.5 | 21.6 ± 8.1 | 23.9 ± 8.1 | .57 | .57 |
| Mood |  |  |  |  |  |
| PANAS positive T1 | 32.0 ± 6.1 | 32.0 ± 5.1 | 32.7 ± 5.5 | .16 | .85 |
| PANAS positive T2 | 26.4 ± 7.7 | 23.9 ± 6.7 | 29.4 ± 8.5 | 3.66 | .03* |
| PANAS negative T1 | 13.2 ± 4.0 | 12.0 ± 2.1 | 12.1 ± 2.2 | 1.54 | .22 |
| PANAS negative T2 | 12.3 ± 3.4 | 11.2 ± 1.8 | 11.1 ± 1.6 | 2.26 | .11 |
| Side effects |  |  |  |  |  |
| Weakness T2 | 1.48 ± 0.63 | 1.45 ± 0.57 | 1.14 ± 0.36 | 3.53 | .03* |
| Dizziness T2 | 1.19 ± 0.40 | 1.52 ± 0.57 | 1.11 ± 0.31 | 6.84 | <.01* |
| Dry mouth T2 | 1.03 ± 0.18 | 1.21 ± 0.41 | 1.04 ± 0.19 | 3.70 | .03* |

BMI = Body Mass Index; MVC = Maximum Voluntary Contraction; PANAS = Positive and Negative Affective Schedule; AQ-k = short version of the Autism Spectrum Quotient; HTAS = Health and Taste Attitudes Questionnaire; STQ = Social Touch Questionnaire; T1 = before drug administration; T2 = ~4.5h after drug administration; M = Mean; SD = Standard Deviation. \* indicates $p < .05$ .

We formulated our hypothesis based on evidence from previous animal research and human studies, which indicates that the hedonic effect associated with the consumption phase solely depends on opioids, while the effect of incentive salience during anticipation also relies on dopamine. Accordingly, we anticipated that participants who received either 400 mg of the dopaminergic antagonist amisulpride or 50 mg of the opioid antagonist naltrexone would exhibit reduced ratings of wanting, effort, and diminished reward-related activity in brain regions associated with incentive salience during reward anticipation (e.g., the ventral tegmental area or VTA, NAc, and vmPFC). On the other hand, we expected that ratings of liking and reward-related activity in brain areas associated with hedonic pleasure during reward consumption (e.g., NAc, OFC) would be reduced in participants receiving 50 mg of the opioid antagonist naltrexone. Furthermore, if social and non-social rewards are subtended by different substrates, dopamine and opioid antagonism should act differently on reward-related brain regions. In particular, based on our previous findings (Korb, Götzendorfer, et al., 2020), we expected stronger pharmacological modulation of food compared to touch rewards, possibly due to the involvement of other neurochemical systems specific to social rewards, such as oxytocin and serotonin (Fischer & Ullsperger, 2017; Tang et al., 2020).

## Results

The statistical analyses described below were pre-registered prior to execution (but after data collection) on Open Science Framework (https://osf.io/6xkph). In each trial of the reward task, participants could receive three levels of social and non-social rewards (high, low, very low; see *Materials and methods* section and Figure 1). The level of reward received depended on which reward level was announced at the beginning of the trial (high or low) and on the force exerted to obtain it (very low rewards were only obtained when participants exerted low effort, which linearly converted into low probability to obtain the announced reward). Differently from Korb et al. (2020), in this study, high, low, and very low reward levels were defined a priori (high = 100% chocolate milk or touch at 6 cm/s, low = 4:1 mix of milk and chocolate milk or touch at 21 cm/s, very low = 100% milk or touch at 27 cm/s). However, the actual wanting and liking of the stimuli did not match those a priori levels for some participants (27,4% for food and 15,7% for touch). Therefore, in contrast to our pre-registration, in both behavioral and fMRI analyses, reward levels (high, low, very low) were re-defined based on the average actual subjective ratings of wanting and liking of each participant.

### No changes in behavioral measures of wanting and liking following opioid and dopamine antagonism

To determine drug effects on the ratings of wanting and liking, and on the effort exerted, we fitted three linear mixed-effects models (LMMs) including Drug (amisulpride, naltrexone, placebo), Reward Type (food, touch), and Reward Level (high, low, very low) as fixed effects, and by-subjects random intercepts and slopes for Reward Type, Reward Level, and their interaction. Null findings were followed up with post hoc Bayesian analysis (see *Materials and methods* section).

The LMM conducted on the ratings of wanting (Figure 2A,C) revealed only a significant main effect of Reward Level (*F*(1, 86.8) = 142.42, *p* < .001). No significant Drug main or interaction effects were observed (all *p* > .14, Figure 2A,C). Bayesian analyses provided moderate support against the Drug effect (BF01 = 6.1) and strong support against the Drug interaction effect (BF01 > 11.7) (see Table S1).

**Figure 2.**
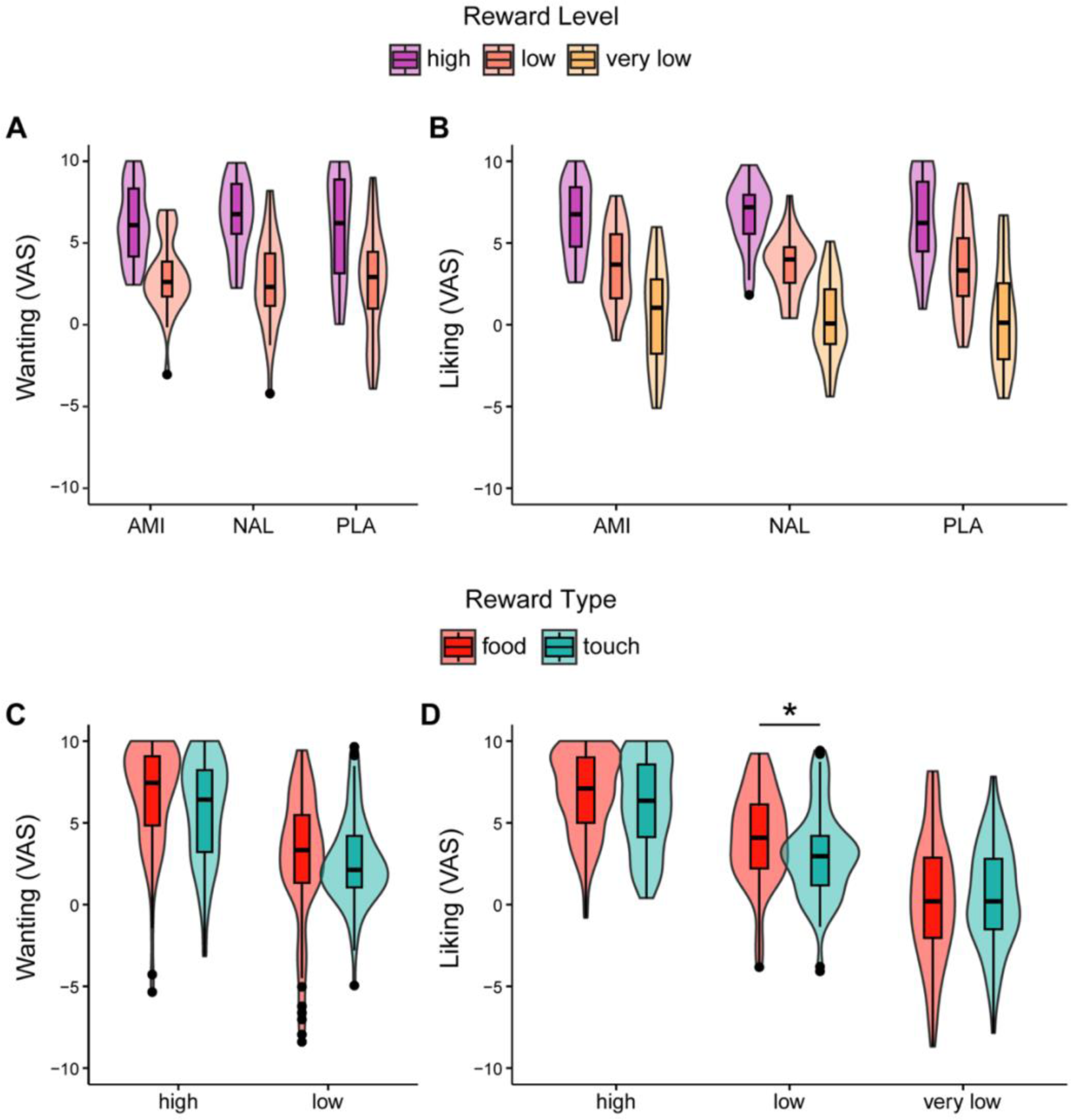
Ratings of Wanting and Liking. No significant drug effects on the ratings of (A) wanting and (B) liking. (C) Ratings of wanting did not differ for food and touch, but (D) ratings of liking were higher for low food rewards compared to low touch rewards (*p* = .04). AMI, amisulpride; NAL, naltrexone; PLA, placebo. * indicates *p* < .05. The violin plots depicted here consist of box plots, representing the median (tick horizontal line), the interquartile range (box), and the lower/upper adjacent values (whiskers) and kernel density plots, representing kernel probability density of the data at different values.

The LMM conducted on the ratings of liking (Figure 2B,D) revealed a significant main effect of Reward Level (*F*(2, 85.5) = 150.00, *p* < .001) and Reward Level by Reward Type interaction (*F*(2, 83.2) = 6.11, *p* < .01). Concerning the interaction, ratings of liking for low non-social rewards were higher than for low social rewards (*p* = .041; Figure 2D). No significant Drug main or interaction effects were observed (all *p* > .34; Figure 2B). Bayesian analyses provided moderate support against the Drug effect (BF01 = 8.4) and strong support against the Drug interaction effects (BF01 > 71.4) (see Table S2).

The LMM conducted on the effort exerted to obtain the announced reward revealed a significant main effect of Reward Level (*F*(1, 86.9) = 68.8, *p* < .001), a Drug by Reward Type interaction (*F*(2, 83.9) = 3.67, *p* = .030), and a Reward Level by Reward Type interaction (*F*(1, 86.1) = 5.92, *p* = .017) (Figure 3). Regarding the Drug by Reward Type interaction (Figure 3B), participants who received amisulpride exerted greater force for non-social compared to social rewards, but the comparison was not statistically significant (*p* = .06, all other comparisons *p* > .35). Regarding the Reward Level by Reward Type interaction (Figure 3C), effort exerted was slightly higher for high non-social compared to high social rewards, but the comparison was not statistically significant (*p* = .15), nor there was a statistically significant difference between the effort exerted for low social and low non-social rewards (*p* = .95). Bayesian analyses provided moderate support against the Drug main effect (BF01 = 4.5, anecdotal BF01 = 1.6 when using a narrower prior), but anecdotal evidence for the Drug by Reward Type interaction effect (BF01 = 1.8) (see Table S3).

**Figure 3.**
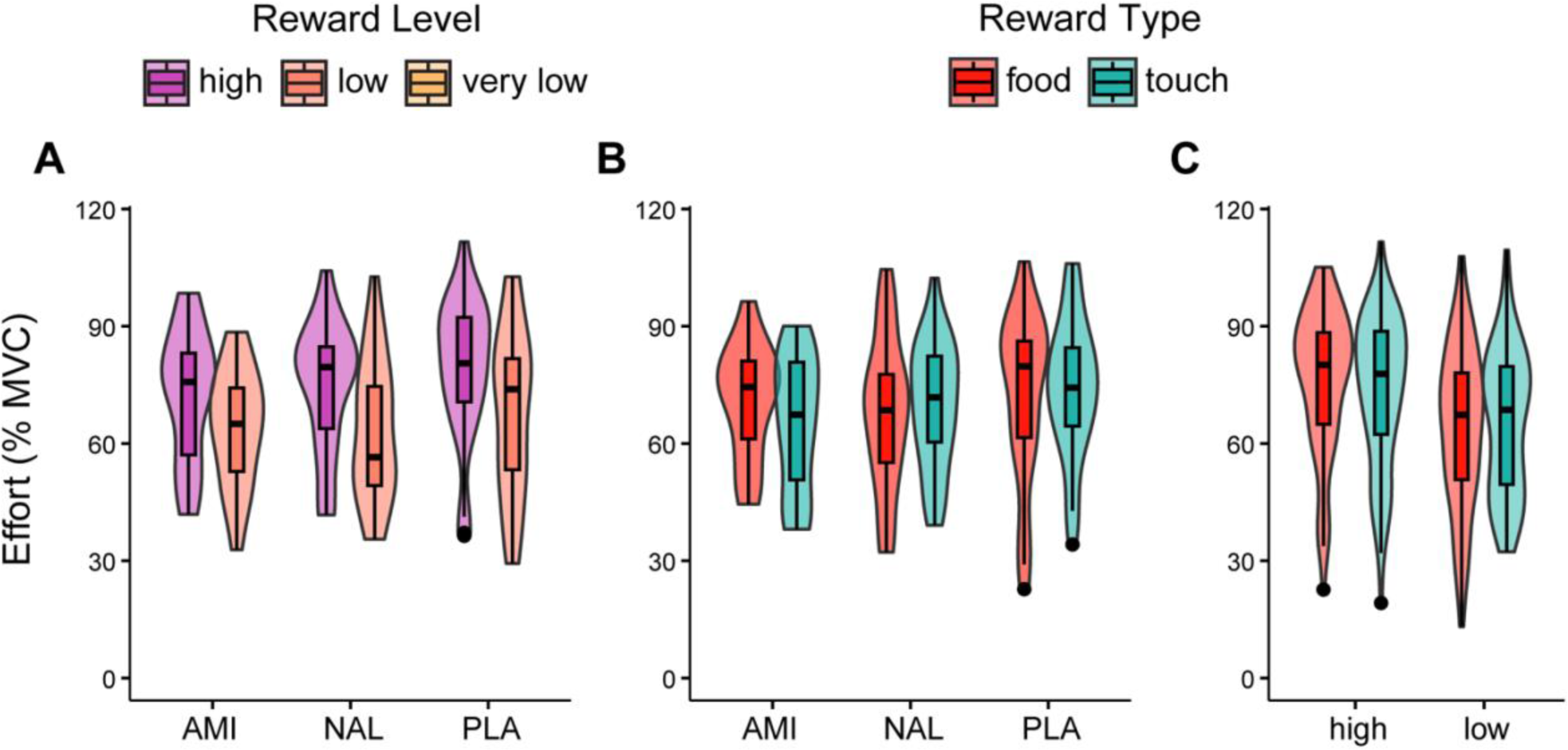
Effort exerted. No significant drug effects on the effort exerted (% of maximum voluntary contraction) to obtain rewards of different (A) level (high, low) and (B) type (food, touch). (C) Effort exerted also did not differ for food and touch independently of the drug group. AMI, amisulpride; NAL, naltrexone; PLA, placebo. The violin plots depicted here consist of box plots, representing the median (tick horizontal line), the interquartile range (box), and the lower/upper adjacent values (whiskers), and kernel density plots, representing kernel probability density of the data at different values.

### Opioid antagonism reduces liking-related activity in the medial orbitofrontal cortex during consumption of social and non-social rewards

Following our pre-registered analysis plan (https://osf.io/6xkph), we conducted whole-brain and region of interest (ROI) analyses, using either the trial-by-trial subjective ratings of wanting, rating of liking, and effort as parametric modulators (GLM.1 and GLM.2), or the categorical predictor Reward Level (high > low, GLM.3). Both GLM.1 and GLM.2 included the ratings of liking as parametric modulator of the delivery phase, and the ratings of wanting and effort as parametric modulators of the anticipation phase. In GLM.1, the ratings of wanting were orthogonalized with respect to effort and therefore included as first and second parametric modulators respectively. In GLM.2 effort was orthogonalized with respect to wanting. For each model, only the first parametric modulator of the anticipation phase was analyzed at the second level (i.e., ratings of wanting for GLM.1 and effort for GLM.2).

Concerning the whole brain analyses, we first investigated the neural correlates of wanting and liking during reward anticipation and consumption, respectively, regardless of reward type and drug group. As expected, reward anticipation was associated with activity in brain regions of the reward system, including the basal ganglia and the prefrontal cortex (Figure 4A). Specifically, linear increase in trial-by-trial ratings of wanting (GLM.1) was associated with increased activation of the pallidum (*t* = 4.75, FWE *p* = .02, peak = [−22 −4 −2]), right lingual gyrus (*t* = 6.85, FWE *p* < .001, peak = [12 −78 −4]), and cerebellum (*z* = 5.71, FWE *p* < .01, peak = [14 −88 −40]) (see also Table S4). Increase in trial-by-trial effort (GLM.2) was associated with increased activity in the medial part of the superior frontal gyrus (*t* = 10.79, FWE *p* < .001, peak = [−4 66 18]), bilateral precentral gyrus (left: *t* = 9.94, FWE *p* < .001, peak = [−38 −20 60]; right: *t* = 7.31, FWE *p* < .001, peak = [38 −18 54]), right middle temporal gyrus (*t* = 7.86, FWE *p* < .001, peak = [60 0 −24]), right middle frontal gyrus (*t* = 7.61, FWE *p* < .001, peak = [30 44 40]), left superior temporal gyrus (*t* = 6.23, FWE *p* < .001, peak = [−44 −24 10]), mid cingulate cortex (*t* = 5.97, FWE *p* = .001, peak = [8 −2 44]) and inferior parietal cortex (*t* = 5.91, FWE *p* = .001, peak = [−34 −60 46]) (complete list in Table S5). Regarding the analysis with the categorical predictor Reward Level (high > low reward; GLM.3), greater BOLD signal during anticipation of high rewards compared to low rewards was observed in the lingual gyrus (*t* = 11.42, FWE *p* < .001, peak = [12 −78 −2]), the medial part of the superior frontal gyrus (*t* = 9.47, FWE *p* < .001, peak = [−6 62 4]), putamen (*t* = 7.66, FWE *p* < .001, peak = [24 2 −8]), bilateral middle temporal gyrus (left: *t* = 9.12, FWE *p* < .001, peak = [−54 −70 22]; right: *t* = 6.43, FWE *p* < .001, peak = [66 −42 8]); right superior temporal gyrus (*t* = 6.11, FWE *p* = .001, peak = 64 −18 −6]), and postcentral gyrus (*t* = 6.33, FWE *p* < .001, peak = [−38 −24 50]) (see Figure 4A and Table S6).

**Figure 4.**
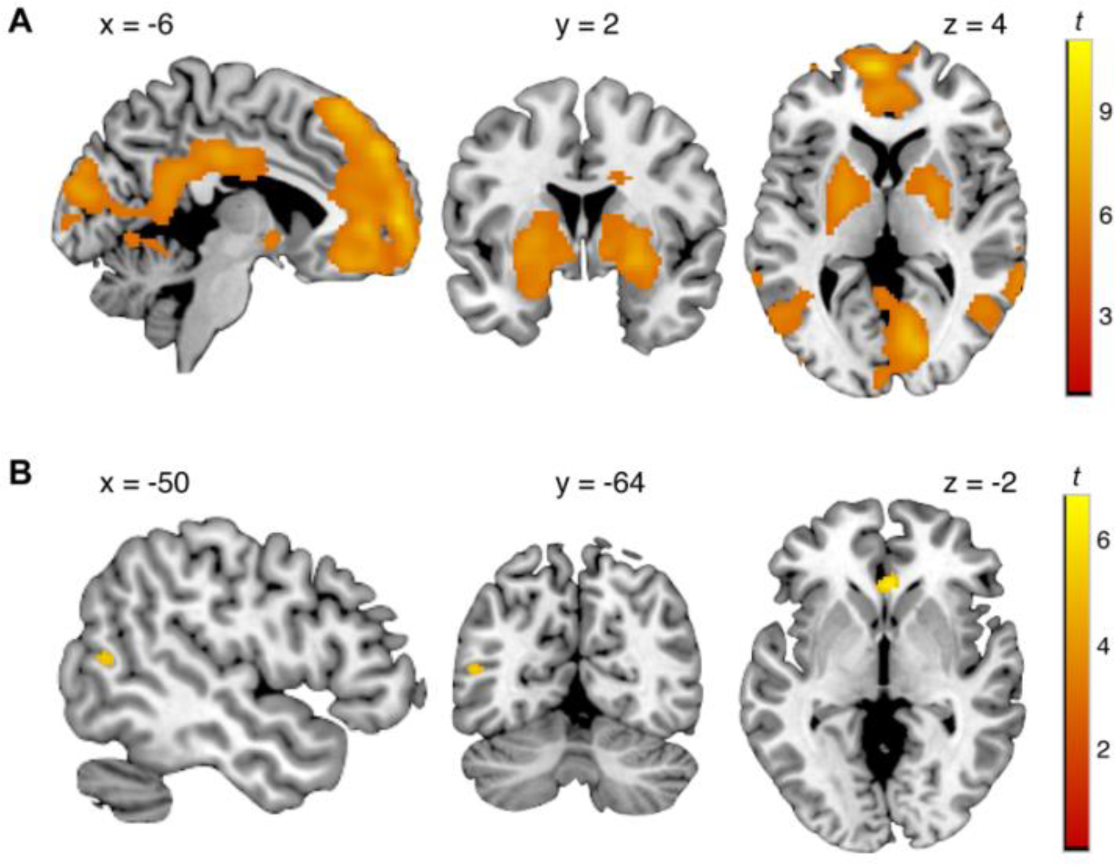
Neural processing during reward anticipation and consumption. Brain regions showing greater activity during high compared to low reward (A) anticipation and (B) consumption (GLM.3). Images are thresholded at *p* < .05 FWE-corrected.

During reward consumption, trial-by-trial ratings of liking did not significantly correlate with the activity in any brain region. In contrast, the analysis with the categorical predictor Reward Level (high > low reward; GLM.3), showed greater activity in the subgenual portion of the anterior cingulate cortex (*t* = 6.64, FWE *p* < .001, peak = [2 28 0]) and left middle temporal gyrus (*t* = 5.23, FWE *p* = .014, peak = [−50 −64 12]) during consumption of high rewards compared to low rewards (Figure 4B, see Table S7).

We did not observe any significant difference between reward types (food and touch) in any of the analyses (see Supplementary Table 4-7).

Regarding drug effects, during reward consumption (GLM.3, high > low), administration of naltrexone was associated with reduced activity compared to placebo in the right medial OFC (*t* = 7.96, FWE *p* < .001, peak = [8 46 −20]; Figure 5A), left medial superior frontal gyrus (dorsomedial prefrontal cortex; *t* = 5.66, FWE *p* < .01, peak = [−6 32 40]), left superior and middle frontal gyrus (*t* = 5.72, FWE *p* < .01, peak = [−16 22 66]; dorsolateral prefrontal cortex: *t* = 5.48, FWE *p* < .01, peak = [−32 34 50]) and right lateral OFC (*t* = 5.32, FWE *p* = .01, peak = [52 40 14]) (see Table S7). Further, during the same phase, administration of amisulpride compared to placebo was associated with reduced activity in the right middle frontal gyrus (premotor cortex; *t* = 7.97, FWE *p* < .001, peak = [44 −4 58]), right thalamus (ventral lateral nucleus; *t* = 5.72, FWE *p* < .01, peak = [14 −16 −2]), left precentral gyrus (primary motor cortex; *t* = 5.59, FWE *p* < .01, peak = [−50 −6 50]), right caudate (*t* = 5.47, FWE *p* < .01, peak = [12 10 14]), Rolandic operculum (*t* = 5.54, FWE *p* < .01, peak = [−62 4 8]) and inferior frontal gyrus (*t* = 5.18, FWE *p* = .02, peak = [−40 32 4]) (see Figure 5B and Table S7). No drug effects were observed during anticipation (GLM.3; Table S6). The analyses including the ratings of wanting and liking and effort exerted as parametric modulators (GLM.1-2) did not reveal any significant drug effects during reward anticipation or consumption (see Table S4 & S5).

**Figure 5.**
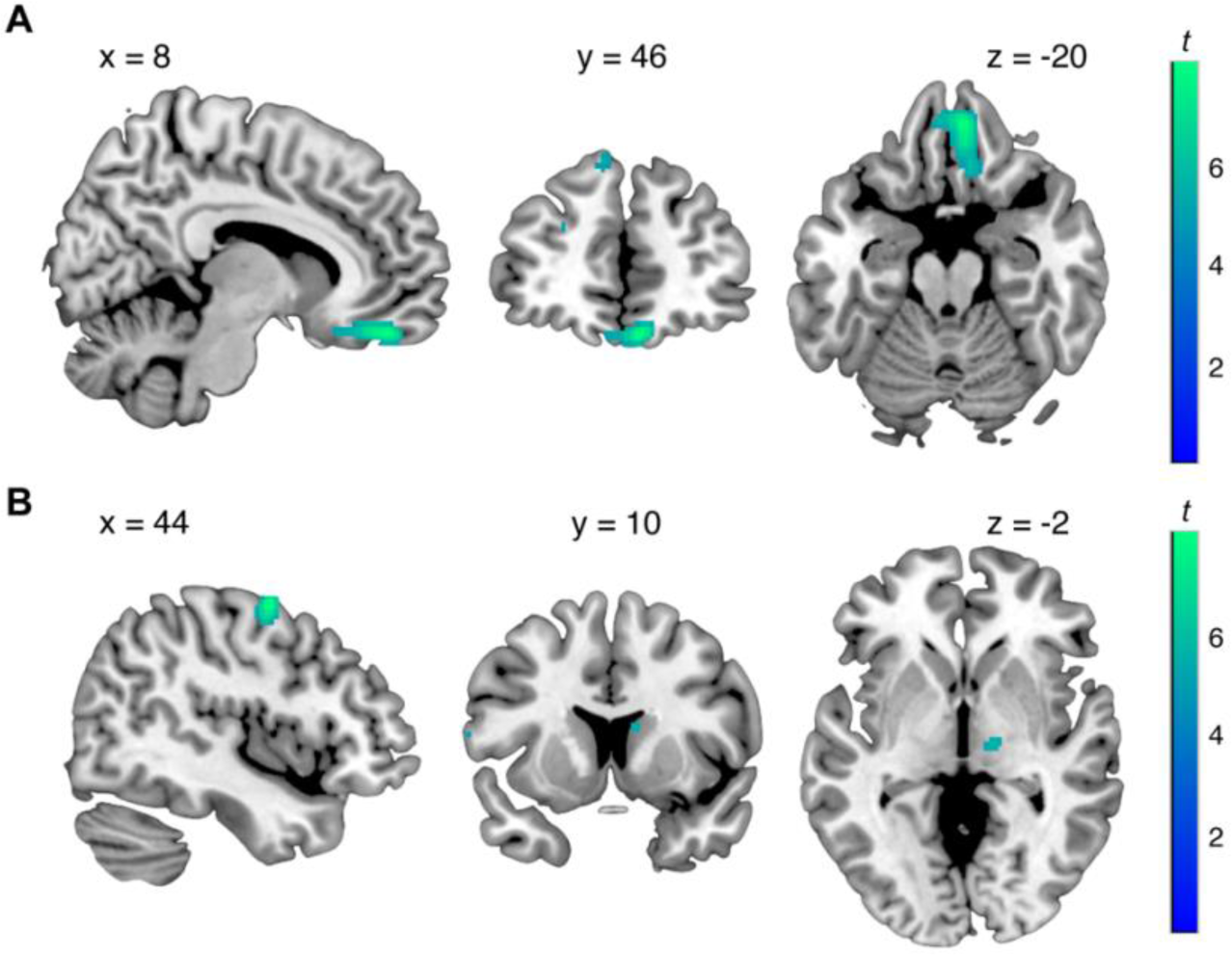
Drug effects on neural processing during reward consumption. (A) Reduced activity in medial orbitofrontal cortex following naltrexone administration compared to placebo (GLM.3). (B) Reduced activity in thalamus, caudate, and middle frontal gyrus following amisulpride administration compared to placebo (GLM.3). Images are thresholded at *p* < .05 FWE-corrected.

In addition to whole brain analyses, we conducted ROI analyses. A priori ROIs were based on anatomical masks and included NAc, VTA, medial OFC, and vmPFC. Brain activity in these ROIs during anticipation and consumption (Phase) of social and non-social rewards (Reward Type) was assessed from the contrasts calculated in GLM.1 (ratings of wanting and liking) and GLM.3 (categorical reward level, high > low) (see *Materials and methods* section for details).

The ROI analysis conducted on the parametrically modulated contrasts in anticipation (wanting ratings) and consumption (liking ratings) (GLM.1) revealed a significant Reward Type x Phase interaction effect (*F*(1,82) = 9.69, *p* = .003) for the activity of the NAc (*F*(1,82) = 10.89, *p* = .001) and medial OFC (*F*(1,82) = 8.61, *p* = .004). Specifically, NAc and medial OFC wanting-related activity was higher during the anticipation of food rewards compared to touch rewards (*p* < .001). Further, for food rewards, NAc and medial OFC wanting-related activity in anticipation was greater than liking-related activity in consumption (*p* < .01). We also observed a significant main effect of Phase for the activity of the vmPFC (*F*(1,82) = 17.34, *p* < .001), reflecting greater wanting-related brain activity during reward anticipation compared to liking-related activity during reward consumption. No significant drug effects were observed. The analysis conducted on the categorical contrasts (high > low reward) in anticipation and consumption (GLM.3) revealed only a significant main effect of Phase for the activity of NAc, VTA, medial OFC, and vmPFC, reflecting a greater increased activity for high compared to low rewards in those regions during anticipation compared to consumption (all *p* < .001).

### Matching of drug groups

Last, we conducted a series of statistical tests to exclude potential group differences in secondary measures, which could have influenced the results.

First, the three drug groups did not significantly differ in terms of age, BMI, maximum voluntary contraction (MVC) before and after the reward task, autistic traits (AQ-k), social touch appreciation (STQ), attitudes to hedonic characteristics of food (HTAS), as well as in their reported mood (PANAS), hunger, and sociability at the beginning of the session (see Table 1 and *Material and methods* section for details). Regarding mood, the naltrexone group reported significantly lower positive mood 4.5 h after pill intake compared to the placebo group (*p* = .03; see Table 1).

Regarding side effects, no significant group differences before drug administration were found. At 4.5 h following drug administration, we observed a significant main effect of Drug for weakness (*F*(2, 85) = 3.53, *p* = .034), dizziness (*F*(2, 85) = 6.84, *p* = .002), and dry mouth (*F*(2, 85) = 3.70, *p* = .029) (Table 1). Participants administered with amisulpride reported greater weakness than participants administered with placebo (*p* = .04). Naltrexone was associated with greater reported dizziness compared to placebo (*p* < .01) and amisulpride (*p* = .02) and greater reported dry mouth compared to amisulpride (*p* = .03). Nevertheless, the average values of all reported side effects were between 1 (“not all”) and 2 (“slightly”), indicating the absence of severe side effects in all drug groups (see Supplementary Figure S1). Drug blinding was successful as participants’ guesses were not significantly related to their group allocation (X^2^(4) = 5.79, *p* = 0.22). Overall, 42% of participants correctly guessed the content of the pill they received.

Finally, no significant differences between drug groups in the average number of high, low, and very low rewards obtained during the task were observed (all *p* > .31, Table 2).

**Table 2.** Average number of high, low, and very low rewards obtained by drug group.

| Reward level | Amisulpride | Naltrexone | Placebo |
| --- | --- | --- | --- |
| High | 35.1% (22.44 ± 5.13) | 36.2% (23.17 ± 9.98) | 37.4% (23.96 ± 8.28) |
| Low | 34.8% (22.25 ± 5.19) | 31.3% (20.00 ± 5.08) | 31.2% (19.96 ± 6.05) |
| Very low | 28.6% (18.31 ± 6.36) | 31.9% (20.43 ± 9.97) | 29.3% (18.75 ± 9.84) |

## Discussion

Prior animal and human findings suggest a partial neurobiological dissociation in the processing of the motivational (wanting) and hedonic (liking) components of reward, with dopaminergic neurotransmission involved in wanting and opioidergic neurotransmission involved in both wanting and liking (see for example Berridge & Kringelbach, 2015; Korb, Götzendorfer, et al., 2020). However, previous studies mainly investigated the neuroanatomical and neurochemical dissociation of reward dimensions within the domain of non-social rewards, such as food or money. Whether a similar neurocircuitry is involved in wanting and liking of rewards of social nature remains unclear. Here, we combined pharmacology and neuroimaging to investigate how blocking of the dopamine and opioid systems modulates the neural processing of wanting and liking of social and non-social rewards. We found that opioid antagonism had an effect at the neural level, where it resulted in reduced activity of the medial orbitofrontal and prefrontal cortices in response to receiving social and non-social rewards. However, contrary to our hypothesis, dopaminergic antagonism did not significantly modulate brain activity related to touch or food anticipation in reward-related regions (e.g., midbrain, NAc), but was instead associated with reduced activity in sensory and motor processing regions during (food and touch) reward consumption. Moreover, we did not observe any effect of the drugs on the behavioral measures of wanting (rating and effort) and liking (rating).

Naltrexone and amisulpride administration did not significantly change subjective wanting and liking of food and touch rewards. Previous research has been inconsistent in regards to the effects of opioid and dopamine antagonism on self-report ratings of wanting and liking, with some studies reporting significant reductions (e.g., Buchel et al., 2018; Soutschek et al., 2021) and others reporting no effects (Korb, Götzendorfer, et al., 2020; Løseth et al., 2019). Especially concerning the effects of dopaminergic manipulations on wanting, most of the effects were observed in studies assessing wanting via more objectively measurable experimental measures, such as effort, rather than subjective ratings (Korb, Götzendorfer, et al., 2020; Soutschek, Gvozdanovic, et al., 2020; Soutschek, Kozak, et al., 2020; Weber et al., 2016). However, in contrast to our previous findings (Korb, Götzendorfer, et al., 2020), we here also failed to show significant changes in the force exerted to obtain the rewarding stimuli following naltrexone and amisulpride administration. This may be due to lower statistical power, as the sample size of this study was tailored mainly to investigate effects on brain activity, rather than on behavior. The post hoc Bayesian analysis provided moderate support for this null finding, but the Bayes factor was smaller than the one found in support for the null effects of drug on the ratings of wanting and liking, and partly depended on prior selection. We therefore refrain from making any claim concerning whether this null finding represent an actual null effect of the pharmacological challenge.

At the neural level, we did not observe dopaminergic modulation of regions of the mesocorticolimbic pathway, usually associated with reward anticipation, such as VTA or NAc. Despite not initially hypothesized, the finding is in line with a recent study that failed to show modulation of wanting-related brain activity for non-consumable goods following the administration of 400 mg of amisulpride (Soutschek et al., 2021). Similarly, using a monetary incentive delay task, Grimm et al. (2021) reported no effects of administration of 200 mg amisulpride and 125 mg L-DOPA on behavioral and neural responses to reward anticipation. Together, the findings highlight the need for a more systematic investigation of dopaminergic modulation of reward processing, which should take into account individual differences (e.g., dopamine baseline levels) and other sources of variation (Martins et al., 2017). We also did not observe any significant effect of blocking opioid receptors on BOLD responses during the anticipation of food and touch stimuli. This is line with a previous study by Büchel et al. (2018) showing that naltrexone reduced neural activity in the mesocorticolimbic system during reward delivery but not during reward anticipation. Similarly, Soutschek et al. (2021) observed a modulation of fronto-striatal connectivity by naltrexone but also reported no significant effects of the opioid antagonist on the neural representations of wanting.

Notably, in the present study, blocking the opioid receptors via naltrexone administration resulted in a reduction of neural activity in the medial OFC during reward consumption. In humans, opioid signaling has been previously linked to hedonic processing of different kinds of rewarding stimuli, such as palatable food (Eikemo et al., 2016; Korb, Götzendorfer, et al., 2020; Nummenmaa et al., 2018), money (Eikemo et al., 2017), or social and erotic stimuli (Buchel et al., 2018; Chelnokova et al., 2014; Hsu et al., 2013; Koepp et al., 2009; Korb, Götzendorfer, et al., 2020). The OFC, a major reward processing hub, is involved in the representation of primary reinforcers, such as taste and touch (Rolls, 2000). Its medial portion, in particular, is involved in encoding reward magnitude during the receipt of different types of reward (Diekhof et al., 2012; Rolls, Cheng, et al., 2020). The OFC receives visual, gustatory, somatosensory, auditory, and olfactory inputs regarding the identity of the stimulus and projects to the striatum, ACC, insula, and inferior frontal gyrus regarding reward value to guide behavior (Kringelbach & Rolls, 2004; Rolls, 2000; Rolls, Cheng, et al., 2020). Previous research robustly showed its involvement in encoding food pleasantness (Kringelbach et al., 2003; Kringelbach & Rolls, 2004; Small et al., 2001).

Importantly, in rodents, opioid stimulation in the OFC causally enhances hedonic reactions to sweet food rewards (Castro & Berridge, 2017). Similar findings were observed in humans. For instance, reduced OFC activation to erotic stimuli was reported following naloxone infusion (Buchel et al., 2018), and opioid availability in the thalamus (measured by PET) was negatively associated with food reward BOLD responses in the OFC (Nummenmaa et al., 2018). The present and previous findings point to an involvement of opioid signaling in the medial OFC related to the consumption of both social and non-social primary rewards, such as food, sex, and touch. However, it is fundamental to remember that our analyses do not allow to distinguish among different neuronal populations in the same brain region. Indeed, distinct but interacting neuronal populations responding to caloric consumption and social interaction have been observed in animal models (Jennings et al., 2019). Further, regardless of the drug group, in the ROI analysis we observed that wanting-related activity in the NAc and medial OFC was mainly elicited by the anticipation of food rewards, but not touch rewards. This is partly in line with previous evidence showing that touch does not consistently result in activity of reward-related brain regions (Sander & Nummenmaa, 2021) and suggests possible differences in neuroanatomical circuitry underlying social and non-social rewards.

Contrary to our initial hypotheses, dopamine antagonism was also associated with neural modulation during reward consumption. Specifically, amisulpride administration reduced activity in sensory and motor processing regions, including the ventrolateral thalamus, pre- and primary motor cortices, and the caudate, compared to placebo. Previous research has shown that these regions are part of the basal ganglia-thalamocortical pathway involved in motor control, also through dopaminergic signaling (Bosch-Bouju et al., 2013). In our study, dopamine antagonism may have affected the processing of the sensorimotor properties of the stimuli (e.g., speed of touch or food taste) rather than reward-related features (Macedo-Lima & Remage-Healey, 2021). However, evidence of such modulation will need further investigation.

Some limitations of the present study should be considered. First, statistical power was on the lower end (also due to drop-outs). Future studies should aim at replicating and extending the present evidence in larger samples or using a within-subject design. Second, despite the long interval between the anticipation and consumption phases in our design, BOLD signal responses related to liking in the consumption phase may have been masked by those pertaining wanting during anticipation, as wanting and liking are highly correlated measures. Future studies should aim to better disentangle these two components by implementing, for example, separated blocks. Additionally, future work should include also an implicit measure of liking, such as facial electromyography (EMG) (Chiappini et al., 2023; Korb et al., 2019). Third, amisulpride can have both presynaptic (increasing dopaminergic transmission) and postsynaptic (blockade of D2/D3 receptors) effects depending on the dosage. The dose of 400 mg employed here is considered the lowest dose to induce postsynaptic effects (Racagni et al., 2004; Schoemaker et al., 1997) and was chosen to ensure the safety of the participants with minimal side effects (as shown by previous studies, e.g., Korb, Götzendorfer, et al., 2020; Soutschek et al., 2021; Weber et al., 2016). Results should be interpreted in the light of the dose employed and of the possible interindividual differences in drug absorption. Finally, it should be noted that modulatory effects of amisulpride and naltrexone were only observed when operationalizing reward-related activity as BOLD responses for high compared to low rewards and not when using the trial-by-trial ratings of wanting/liking and effort exerted as parametric modulators. This may be due to a lack of power for such parametric analysis and individual variability in these measures. Null drug findings were also observed for the ROI analyses. Therefore, these results should be interpreted cautiously and must be replicated in a larger sample.

In conclusion, by combining pharmacological challenges, neuroimaging, and a behavioral paradigm inspired by animal research, we showed that blocking the opioid system reduced liking-related activity in OFC during the receipt of primary social and non-social stimuli. The findings represent a significant first step in deepening our understanding of the neurochemical and neuroanatomical foundations of wanting and liking of social and non-social rewards. This research holds promise for enhancing our comprehension of psychopathologies characterized by general disturbances in reward processing, such as anorexia nervosa and depression.

## Materials and methods

### Sample

Ninety-four healthy volunteers (57 females) aged 18-35 took part in the study. Five participants did not undergo MR scanning because of sickness (3), claustrophobia (1), or impossibility to provide a urine sample for the drug/pregnancy test (1) and were therefore excluded, leading to a final sample size of 89 (age *M* = 23.7, *SD* = 3.7; but N = 85 for fMRI data analyses, see below). All participants reported being right-handed, to smoke less than five cigarettes daily, to have no history of current or former drug abuse, to like milk and chocolate, not to suffer from diabetes, lactose intolerance, lesions, or skin diseases on the left forearm, and to be free of psychiatric or neurological disorders. Other exclusion criteria included contraindications to MRI (e.g., metallic implants) and having taken part in a pharmacological study in the 2 months before the testing appointment. To avoid sexual connotations of the social touch, the tactile stimulation was always administered by a same-gender experimenter, and only participants who reported to be heterosexual were included. Person-related variables included age, BMI, autistic traits (Short version of the German Autism Spectrum Quotient, AQ-k; Freitag et al., 2007), social touch appreciation (Social Touch Questionnaire, STQ; Wilhelm et al., 2001), taste attitudes (subscales “craving for sweet foods”, “using food as a reward”, “pleasure” of the Health and Taste Attitudes Questionnaire, HTAS; Roininen & Tuorila, 1999), hunger, and sociability before the reward task (Table 1).

The study was approved by the Ethical Committee of the Medical University of Vienna (EK N. 1393/2017) and was performed in line with the Declaration of Helsinki (World Medical Association, 2013). Participants signed the informed consent and received a monetary compensation of 90€.

### Stimuli

In the non-social condition, chocolate milk, a 4:1 mix of milk and chocolate milk, and milk were used respectively as high, low, and very low reward. The three stimuli were identical in fat and sugar content (1.5 g of fat, 10 g of sugar per 100 g). Tap water was used to rinse the mouth at the end of each trial. The initial stimulus temperature of these liquids was kept constant (∼4° C) across participants. Stimuli were delivered by means of computer-controlled pumps (PHD Ultra pumps, Harvard Apparatus) attached to plastic tubes (internal ø 1,6 mm; external ø 3,2 mm; Tygon tubing, U.S. Plastic Corp.), which ended jointly in the participants’ mouth. In each trial, 1.5 mL of liquid was administered for 2 s. Overall, including practice trials, participants consumed 102 ml of liquids, composed of 51 ml of water, and 51 ml of sweet milk with different concentrations of chocolate aroma (depending on effort, see below).

In the social condition, rewards consisted of gentle caresses over a marked nine-cm area of the participants’ right forearm. A same-gender experimenter applied three different caressing frequencies (six cm/s, 21 cm/s, 27 cm/s) for six seconds, constituting the high, low, and very low social rewards, respectively. To facilitate stroking, the experimenter received extensive training and was guided during each trial of the task by a sound indicating the rhythm for stimulation, heard via headphones.

### Procedure

A randomized, double-blind, placebo-controlled, three-armed study design was implemented. Participants came to the laboratory twice, with a maximum interval of 2 months between the two appointments. At the first visit (T0), participants received a health screening, including ECG and blood examination, as well as a neuropsychiatric interview using the Mini-International Neuropsychiatric interview (M.I.N.I). The described experiment was carried out at the second visit (T1). To enhance and equate drugs’ absorption time, participants were instructed not to eat in the preceding 6 hours before coming to T1. At the arrival, they filled out the Positive and Negative Affect Scale (PANAS; Watson et al., 1988) and a questionnaire about physical symptoms (see Supplementary Figure S1), performed a urine drug test sensitive to opiates, amphetamine, methamphetamine, cocaine, and other substances, and, if females, a urine pregnancy test. If the drug and pregnancy tests resulted negative, participants received a capsule filled with either 400 mg of amisulpride (Solian®; N = 32), 50 mg of naltrexone (Dependex®; N = 29), or 650 mg of mannitol (sugar; N = 28) from the study doctor. Following drug intake, participants received a snack and, after a waiting time of 3 hours, they were transferred to the MRI center.

Participants first experienced each social and non-social reward once outside the MR-scanner. After being installed in the MR-scanner, the participants’ MVC was established. Participants were required to squeeze the dynamometer with their left hand as hard as possible for three seconds, three times. The peak force (in Newton) across the three trials represented participants’ MVC, used as threshold for the effort task. Following calibration, participants performed two training trials of the social and non-social conditions to familiarize themselves with the procedure.

Participants performed in total four blocks of the reward task (acquired in four fMRI runs) (Chiappini et al., 2022), two in the social and two in the non-social condition. The order of the blocks (ABAB or BABA) was randomized across participants. Each block included 16 trials, including the following main components: i) announcement of reward (high or low; 1s); ii) rating of wanting on a visual analog scale (VAS) from −10 (not at all) to +10 (very much) (4s); iii) effort task: to obtain the announced reward, participants had to squeeze the hand dynamometer (the applied force was translated into a probability of obtaining the reward; 4s); iv) announcement of reward gained (high, low, or, in case of low effort, very low; 1s); v) prepare for reward delivery (6s); vi) reward delivery (7s); vii) relax phase (8s); viii) rating of liking on VAS from −10 (not at all) to +10 (very much) (4s); ix) only in the non-social condition, water delivery for rinsing.

After completing the task, the participants’ MVC was assessed again. Then participants completed the PANAS and physical symptoms questionnaire (∼4.5 h after drug intake). Approximately 5.5 h after drug intake, a blood sample was collected. At the end of the session, participants were asked to guess the identity of the drug they received and were debriefed about the aim of the study.

### Drug administration

Naltrexone is a non-selective opioid antagonist with high affinity to the µ- and κ-opioid receptors, while amisulpride is a selective dopamine D2/D3 receptor antagonist. We used 50 mg per-oral naltrexone (Dependex®), which blocks more than 90% of µ-opioid receptors (Lee et al., 1988), and 400 mg per-oral amisulpride (Solian®), which results in 50–80% D2 receptor blockade (la Fougère et al., 2005; Meisenzahl et al., 2008). Higher doses of amisulpride are not recommended in research on healthy subjects due to the increased risk of extrapyramidal side effects. The length of the time interval between drug intake and task onset (3 h) was modeled on previous studies using the same compounds and doses (Korb, Götzendorfer, et al., 2020; Weber et al., 2016) and on drugs’ pharmacodynamics. Amisulpride reaches a first peak in serum after 1 h and a second (higher) peak after approximately 4 h, with a 12 h half-life (Rosenzweig et al., 2002). Naltrexone reaches maximal concentration in plasma after 1 h and has a half-life of approximately 4 h (Meyer et al., 1984).

### MRI acquisition and data pre-processing

MRI data were acquired using a 3T Siemens Prisma fit MRI scanner (Siemens Healthineers, Erlangen, Germany) with a 64-channel head coil. Functional whole-brain scans were collected using a multiband-accelerated T2*-weighted 2D echoplanar imaging (EPI) sequence (40 slices, TE/TR = 35/1000 ms, flip angle = 62°, voxel size = 2.3 × 2.3 × 3.0 mm, FOV = 220 x 220 mm). Structural images were acquired with a magnetization-prepared two inversion time rapid gradient-echo (MP2RAGE) sequence (TE/TR = 2.98/4000ms, flip angle =4°, voxel size = 1 × 1 × 1 mm, FOV = 256 x 216 x 160 mm). Imaging data were pre-processed with Statistical Parametric Mapping (SPM12; Wellcome Trust Centre for Neuroimaging, London, UK) and FMRIB Software Library (FSL; Analysis Group, FMRIB, Oxford, UK). Pre-processing included: realignment to the first image of each run, magnetic field inhomogeneity distortion correction (topup; Andersson et al., 2003), co-registration to T1 image, segmentation, normalization to MNI template space, smoothing with an 8 mm full width at half-maximum (FWHM) Gaussian kernel. Runs containing framewise displacement greater than 0.5 mm on more than 40% of the total frames were excluded from additional analyses (1 food run in 6 participants). Following smoothing, the ICA-AROMA algorithm was applied to reduce motion artifacts (Pruim et al., 2015). Due to technical issues, fMRI data of one participant is not available and data from three participants could not be analyzed as they completed only the social touch blocks. fMRI analyses thus included 85 participants in total (30 AMI, 27 NAL, 27 PLA).

## Statistical analyses

### Behavioral data

Statistical analyses of behavioral data were conducted in R (R Core Team, 2021). The effect of the administered drug on wanting, liking, and effort for social and non-social rewards was investigated using three LMMs (one for each behavioral dependent variable). Each model included Drug (amisulpride, naltrexone, placebo), Reward Type (food, touch), and Reward Level (high, low, very low) as fixed effects, and by-subjects random intercepts and slopes for Reward Type, Reward Level, and their interaction. Group comparisons for age, scores of the trait questionnaires, BMI, and MVC were assessed with analysis of variance (ANOVA). Planned comparisons were corrected using the Tukey method.

#### Post hoc Bayesian analyses

Null findings in behavioral analyses were followed up with Bayesian analyses in JASP 0.15 (JASP team, 2021). We implemented repeated measures ANOVAs with a default multivariate Cauchy prior (r scale prior width for fixed effects =.5). Then, robustness checks were conducted using a narrower (r = .2) and a wider prior (r = 1), as recommended by Van Doorn et al. (2021). We computed Bayes factors (BF01) to estimate the evidence in favor of both the null (H0) and the alternative hypotheses (H1, drug model), using the following thresholds: moderate support for H0 with a BF01 between 3 and 10, strong support for H0 with a BF01 larger than 10, moderate support for H1 with BF01 between 0.3 and 0.1 and strong support for H1 with a BF01 smaller than 0.1 (van Doorn et al., 2021).

### fMRI data

#### Whole brain analysis with parametric modulation

Pre-processed data were analyzed as an event-related design in the context of the GLM approach in a two-level procedure. At the first level, two design matrices, one for food runs and one for touch runs, were fitted for each subject. Each design matrix included regressors for each distinct phase of the trials (fix cross, anticipation pre-effort, rate wanting, effort, anticipation post-effort, prepare delivery, delivery, relax, rate liking, prepare rinsing (only for food trials), rinsing (only for food trials)). To explore the brain activation in response to wanting during reward anticipation (anticipation pre-effort) and to liking during and after reward consumption (delivery), parametric modulation was implemented at the first level. Two models were fitted with each of the two parametric modulators for the anticipation phase (ratings of wanting and effort exerted). In the first model (GLM.1), the ratings of wanting were orthogonalized with respect to effort and therefore included as first and second parametric modulators of the anticipation pre-effort phase, while the ratings of liking were included as parametric modulator of the delivery phase. The second model (GLM.2) was identical to the first one, but, for the phases of anticipation pre-effort, effort was orthogonalized with respect to wanting. We further analyzed only the first parametric modulator for each model, ignoring the second one. For each design matrix (food runs and touch runs), the following contrast images were calculated and taken to the second-level analysis: (i) first-order parametric modulation of ratings of wanting (GLM.1) or effort (GLM.2) in anticipation pre-effort, and (ii) first-order parametric modulation of ratings of liking in delivery.

Second-level analysis included the factors Drug (amisulpride, naltrexone, placebo) and Reward Type (food, touch). A mixed-model ANOVA (flexible factorial design) was fitted for each first-level contrast to explore the main effects of Drug and Reward Type, as well as their interaction.

#### Whole brain analysis with categorical predictor Reward Level (high > low)

In order to reproduce the analysis performed in the majority of the previous papers (e.g., Buchel et al., 2018; Rademacher et al., 2010; Spreckelmeyer et al., 2009), another first-level analysis (GLM.3), including two design matrices for food runs and touch runs separately, were fitted for each participant, using the categorical variable Reward Level (high > low reward) instead of the continuous parametric modulators (trial by trial ratings of wanting and liking, and effort exerted). The category “low reward” included both low and very low reward types. The simple contrasts of high reward vs low reward in the main phases of (i) reward anticipation (anticipation pre-effort) and (ii) reward consumption (delivery) were taken to the second-level for group comparison. Similar to what was explained above, a mixed-model ANOVA (flexible factorial design) was fitted for each phase (anticipation pre-effort, delivery) to explore the main effects of Drug and Reward Type, as well as their interactions. All reported results are based on family-wise error (FWE) correction for voxel intensity tests (FEW *p* < .05).

#### ROI analyses

In addition to the whole-brain analyses, we performed a region of interest (ROI) analysis with a priori defined masks. ROIs were chosen based on findings from previous metanalyses on monetary, erotic, food, and social rewards (Gu et al., 2019; Sescousse et al., 2013) and relevant neuroimaging research (e.g., Izuma et al., 2008; O’Doherty et al., 2002; Rademacher et al., 2010) and include 4 ROIs generated based on anatomical masks: NAc (AAL3 atlas), VTA (Trutti et al., 2021), medial OFC (Jülich Brain MPM atlas) and vmPFC (AAL3 atlas). For each subject, brain activity in each ROI was extracted for the following contrast images: (i) first-order parametric modulation of rating of wanting in anticipation pre-effort and (ii) first-order parametric modulation of rating of liking in delivery. To investigate the effect of the administered drugs on social and non-social reward anticipation and consumption, ANOVAs (one for each ROI) were performed, including the between-subject factor Drug (amisulpride, naltrexone, placebo), and the within-subject factors Reward Type (food, touch) and Phase (anticipation, consumption).

The same ROI analysis was conducted also on the contrasts calculated from GLM.3: high > low reward in i) anticipation and ii) consumption. We adjusted the threshold of statistical significance for both ROI analyses based on the number of ROIs (*p* < .0125).

## Data availability

The behavioral data and analysis scripts that support the findings of this data are available on Open Science Framework https://osf.io/kw623/.

## Author contributions (CRediT)

Conceptualization: GS, CE, SK, MW; Methodology: GS, SK, JL, CW; Software: SK; Formal analysis: CM; Visualization: CM; Investigation: SG, EC; Data curation: CM, SK, EC, SG; Supervision: GS, SK; Project administration: SK, MW, EC, SG; Funding acquisition: GS, CE; Writing – original draft: CM; Writing – review and editing: CM, GS, SK, JL, EC, CW

## Acknowledgments

This work was supported by the Vienna Science and Technology Fund (WWTF) with a grant (CS15-003) awarded to GS and CE. The funding sources had no role in the elaboration of the study design, the data collection, analysis and interpretation, the writing of this report, and the decision to submit this manuscript for publication. We thank Eva Pool for valuable input regarding the fMRI analyses, Gheorghe L. Preda for his contribution in carrying out the medical procedures, and the students involved in data collection: Raimund Buehler, Merit Pruin, Björn Bartuska, Luca Wiltgen, Ariane Hohl, Berit Hansen.

## Supplementary Material

**Figure S1.**
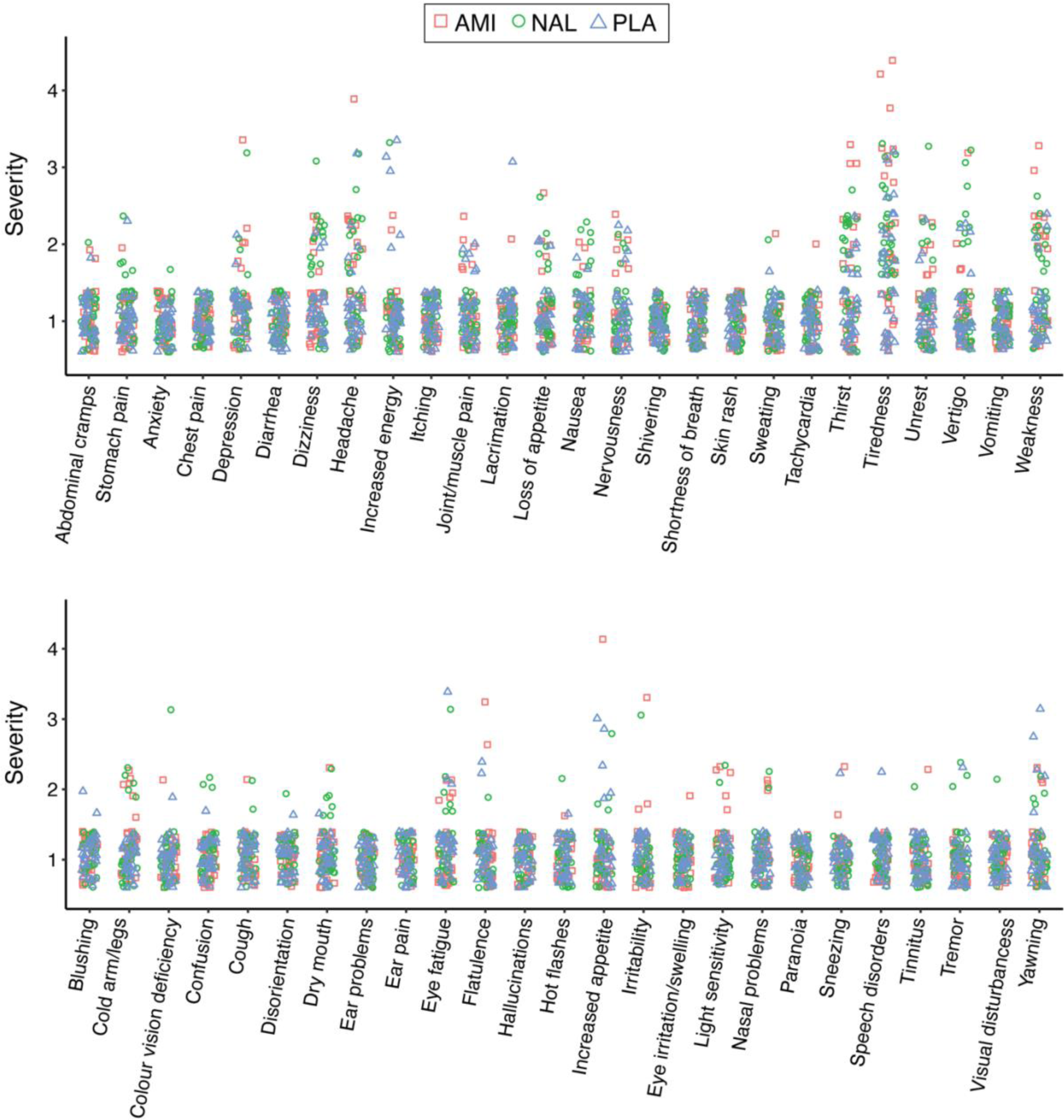
Individual scores of reported side effects assessed 4.5 h following amisulpride (AMI), naltrexone (NAL), and placebo (PLA) administration. Possible side effects were measured on a 4-point Likert scale (1 = “not at all”, 2 = “slightly”, 3 =“moderately”, and 4 =“very much“) immediately before and 4.5 h after drug administration. The list of side effects (original in German) was: headache; unrest; nervousness; stomach ache; abdominal cramps; nausea; vomiting; joint/muscle pain; weakness; anxiety; thirst; swindle; dizziness; chills; sweating; tiredness; lacrimation; tachycardia; chest pain; shortness of breath; diarrhea; skin rash; itching; loss of appetite; increased energy; depressed; irritability; hallucinations; confusion; paranoia; disorientation; tremor; eye irritation/swelling; light sensitivity; eye pain/fatigue; color vision deficiency; ear problems; ear pain; blushing; nasal problems; sneezing; coughing; increased yawning; flatulence; dry mouth; cold arms/legs; hot flashes; increased appetite; visual disturbances; speech disorders; tinnitus.

**Table S1.** Results of the Bayesian rmANOVA on the ratings of wanting.

| Models | BF01 (error %) |  |  |
| --- | --- | --- | --- |
|  | default r | r = 1 | r = .2 |
| Null model (incl. type, level, type*level, subject, and random slopes) | 1.0 | 1.0 | 1.0 |
| drug | 6.1 (5.9) | 18.0 (5.1) | 3.7 (19.8) |
| drug + drug*type | 11.7 (4.6) | 81.4 (4.3) | 3.0 (22.1) |
| drug + drug*level | 35.7 (4.7) | 309.3 (4.3) | 6.2 (23.9) |
| drug + drug*type + drug*level | 64.5 (5.2) | 1312.3 (4.8) | 5.4 (29.8) |
| drug + drug*type + drug*level + drug*type*level | 146.2 (58.0) | 9893.2 (48.4) | 8.4 (37.2) |

Note. All models include type, level, type\*level, subject, and random slopes for all repeated measures factors.

**Table S2.** Results of the Bayesian rmANOVA on the ratings of liking.

| Models | BF01 (error %) |  |  |
| --- | --- | --- | --- |
|  | default r | r = 1 | r = .2 |
| Null model (incl. type, level, type*level, subject, and random slopes) | 1.0 | 1.0 | 1.0 |
| drug | 8.4 (20.9) | 21.1 (4.0) | 4.0 (31.2) |
| drug + drug*type | 71.4 (23.1) | 429.7 (3.6) | 6.2 (26.4) |
| drug + drug*level | 331.8 (30.0) | 5255.5 (3.6) | 11.2 (30.5) |
| drug + drug*type + drug*level | 1440.8 (42.2) | 102443.8 (3.7) | 45.5 (33.1) |
| drug + drug*type + drug*level + drug*type*level | 35529.9 (25.3) | 3.047x10<sup>+6</sup> (40.7) | 100.1 (41.1) |

Note. All models include type, level, type\*level, subject, and random slopes for all repeated measures factors.

**Table S3.** Results of the Bayesian rmANOVA on the effort exerted to obtain the reward.

| Models | BF01 (error %) |  |  |
| --- | --- | --- | --- |
|  | default r | r = 1 | r = .2 |
| Null model (incl. type, level, type*level, subject, and random slopes) | 1.0 | 1.0 | 1.0 |
| drug | 4.5 (20.7) | 5.8 (6.0) | 1.6 (4.3) |
| drug + drug*type | 1.8 (23.1) | 6.3 (4.7) | 0.7 (5.3) |
| drug + drug*level | 13.0 (23.6) | 115.3 (5.1) | 4.3 (24.0) |
| drug + drug*type + drug*level | 19.5 (28.3) | 117.0 (4.8) | 1.7 (22.3) |
| drug + drug*type + drug*level + drug*type*level | 89.5 (35.3) | 1348.1 (64.3) | 4.2 (27.2) |

Note. All models include type, level, type\*level, subject, and random slopes for all repeated measures factors.

**Table S4.** fMRI whole brain results during reward anticipation (parametric modulation of the ratings of wanting, GLM.1).

|  | Hemi | k | T | Z | P<sub>FWE</sub> | x | y | z |
| --- | --- | --- | --- | --- | --- | --- | --- | --- |
| *Linear increase Wanting* |  |  |  |  |  |  |  |  |
| **Lingual gyrus** | R | **269** | **6.85** | **6.08** | **.000** | **12** | **-78** | **-4** |
| Cerebellum | R |  | 5.55 | 5.10 | .004 | 12 | -74 | -18 |
| **Cerebellum** | R | **15** | **5.71** | **5.23** | **.002** | **14** | **-88** | **-40** |
| **Pallidum** | R | **16** | **5.11** | **4.75** | **.021** | **-22** | **-4** | **-2** |
| *Food > Touch* |  |  |  |  |  |  |  |  |
| No suprathreshold clusters |  |  |  |  |  |  |  |  |
| *Touch > Food* |  |  |  |  |  |  |  |  |
| No suprathreshold clusters |  |  |  |  |  |  |  |  |
| *PLA > AMI* |  |  |  |  |  |  |  |  |
| No suprathreshold clusters |  |  |  |  |  |  |  |  |
| *PLA > NAL* |  |  |  |  |  |  |  |  |
| No suprathreshold clusters |  |  |  |  |  |  |  |  |
| *PLA (food >touch) > AMI (food > touch)* |  |  |  |  |  |  |  |  |
| No suprathreshold clusters |  |  |  |  |  |  |  |  |
| *PLA (food >touch) > NAL (food > touch)* |  |  |  |  |  |  |  |  |
| No suprathreshold clusters |  |  |  |  |  |  |  |  |

Note. Hemi, hemisphere; cluster size, k (extended threshold k = 15); x y z, MNI coordinates; Brain regions were labelled with the Automated Anatomical Labeling Atlas (AAL) 3 (Rolls, Huang, et al., 2020).

**Table S5.** fMRI whole brain results during reward anticipation (parametric modulation of the effort exerted to obtain the rewards; GLM.2)

|  | Hemi | k | T | Z | P<sub>FWE</sub> | x | y | z |
| --- | --- | --- | --- | --- | --- | --- | --- | --- |
| *Linear increase Effort* |  |  |  |  |  |  |  |  |
| **Superior frontal gyrus medial** | L | **39019** | **10.79** | **Inf** | **.000** | **-4** | **66** | **18** |
| Superior frontal gyrus medial | L |  | 10.60 | Inf | .000 | -4 | 58 | 34 |
| Middle occipital gyrus | L |  | 10.14 | Inf | .000 | -22 | -98 | 16 |
| **Precentral gyrus** | L | **2478** | **9.94** | **Inf** | **.000** | **-38** | **-20** | **60** |
| Precentral gyrus | L |  | 6.58 | 5.88 | .000 | -18 | -18 | 70 |
| Supramarginal gyrus | L |  | 6.57 | 5.87 | .000 | -52 | -26 | 30 |
| **Middle temporal gyrus** | R | **521** | **7.86** | **6.77** | **.000** | **60** | **0** | **-24** |
| Temporal pole | R |  | 7.15 | 6.29 | .000 | 54 | 16 | -28 |
| Inferior temporal gyrus | R |  | 5.63 | 5.16 | .003 | 56 | 6 | -36 |
| **Precentral gyrus** | R | **1394** | **7.31** | **6.39** | **.000** | **38** | **-18** | **54** |
| Precentral gyrus | R |  | 6.75 | 6.00 | .000 | 36 | -20 | 42 |
| Precentral gyrus | R |  | 5.04 | 4.69 | .023 | 48 | -6 | 60 |
| **Middle frontal gyrus** | R | **91** | **7.16** | **6.29** | **.000** | **30** | **44** | **40** |
| **Cerebellum** | R | **118** | **6.41** | **5.75** | **.000** | **24** | **-76** | **-54** |
| Cerebellum crus | R |  | 5.34 | 4.93 | .008 | 36 | -80 | -48 |
| **Cuneus** | R | **66** | **6.28** | **5.66** | **.000** | **18** | **-82** | **42** |
| **Superior temporal gyrus** | L | **182** | **6.23** | **5.62** | **.000** | **-44** | **-24** | **10** |
| Superior temporal gyrus | R |  | 5.71 | 5.23 | .002 | -54 | -20 | 6 |
| **Middle cingulate cortex** | L | **221** | **5.97** | **5.42** | **.001** | **8** | **-2** | **44** |
| Middle cingulate cortex | R |  | 5.38 | 4.96 | .007 | -2 | -6 | 50 |
| **Inferior parietal cortex** | L | **84** | **5.91** | **5.38** | **.001** | **-34** | **-60** | **46** |
| **Middle temporal gyrus** | R | **142** | **5.72** | **5.23** | **.002** | **64** | **-36** | **6** |
| **Precentral gyrus** | L | **39** | **5.71** | **5.22** | **.002** | **-44** | **8** | **46** |
| **Temporal pole** | R | **30** | **5.62** | **5.16** | **.003** | **34** | **12** | **-40** |
| **Cerebellum** | L | **40** | **5.49** | **5.05** | **.005** | **-2** | **-54** | **-2** |
| **Heschl gyrus** | R | **27** | **5.37** | **4.96** | **.007** | **42** | **-18** | **12** |
| **Middle frontal gyrus** | R | **27** | **5.24** | **4.85** | **.012** | **48** | **36** | **28** |
| *Food > Touch* |  |  |  |  |  |  |  |  |
| No suprathreshold clusters |  |  |  |  |  |  |  |  |
| *Touch > Food* |  |  |  |  |  |  |  |  |
| No suprathreshold clusters |  |  |  |  |  |  |  |  |
| *PLA > AMI* |  |  |  |  |  |  |  |  |
| No suprathreshold clusters |  |  |  |  |  |  |  |  |
| *PLA > NAL* |  |  |  |  |  |  |  |  |
| No suprathreshold clusters |  |  |  |  |  |  |  |  |
| *PLA (food > touch) > AMI (food > touch)* |  |  |  |  |  |  |  |  |
| No suprathreshold clusters |  |  |  |  |  |  |  |  |
| *PLA (food > touch) > NAL (food > touch)* |  |  |  |  |  |  |  |  |
| No suprathreshold clusters |  |  |  |  |  |  |  |  |

Note. Hemi, hemisphere; cluster size, k (extended threshold k = 15); x y z, MNI coordinates; PLA, placebo; AMI, amisulpride; NAL, naltrexone. Brain regions were labelled with the Automated Anatomical Labeling Atlas (AAL) 3 (Rolls, Huang, et al., 2020).

**Table S6.**
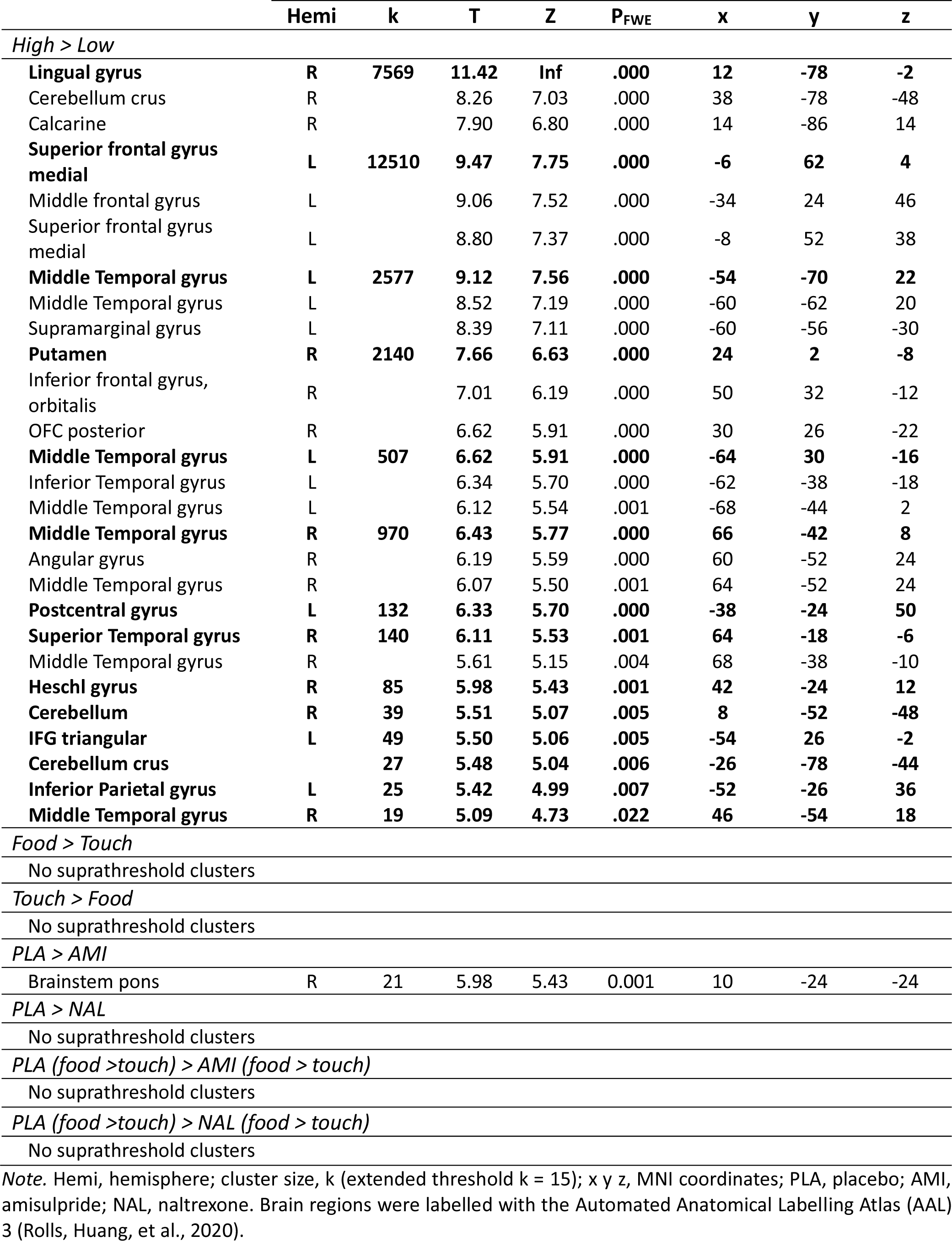
fMRI whole brain results during reward anticipation (High reward > Low reward; GLM.3)

**Table S7.**
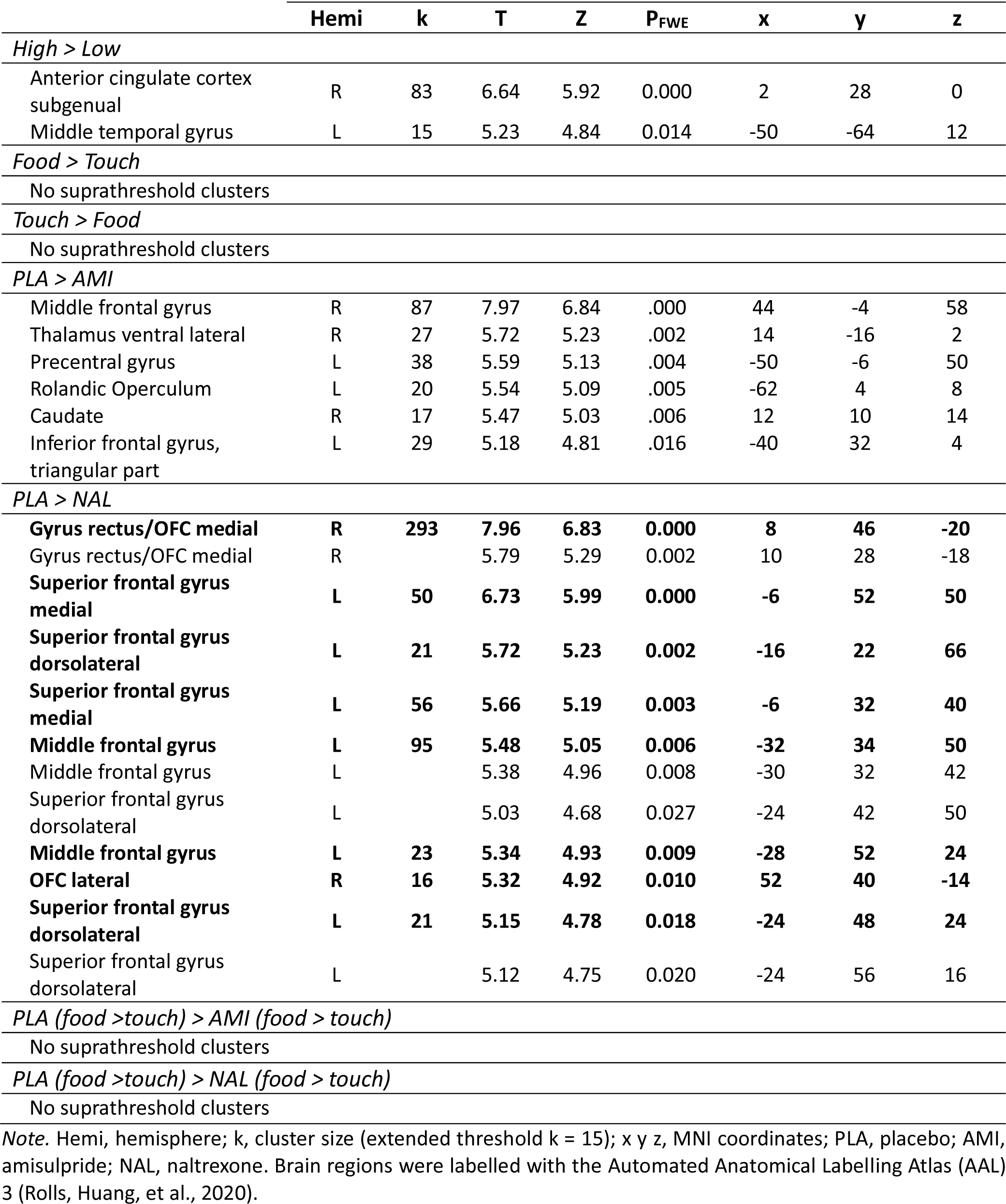
fMRI whole brain reksults during reward consumption (High reward > Low reward; GLM.3)

|  | Hemi | k | T | Z | P<sub>FWE</sub> | x | y | z |
| --- | --- | --- | --- | --- | --- | --- | --- | --- |
| *High > Low* |  |  |  |  |  |  |  |  |
| Anterior cingulate cortex subgenual | R | 83 | 6.64 | 5.92 | 0.000 | 2 | 28 | 0 |
| Middle temporal gyrus | L | 15 | 5.23 | 4.84 | 0.014 | -50 | -64 | 12 |
| *Food > Touch* |  |  |  |  |  |  |  |  |
| No suprathreshold clusters |  |  |  |  |  |  |  |  |
| *Touch > Food* |  |  |  |  |  |  |  |  |
| No suprathreshold clusters |  |  |  |  |  |  |  |  |
| *PLA > AMI* |  |  |  |  |  |  |  |  |
| Middle frontal gyrus | R | 87 | 7.97 | 6.84 | .000 | 44 | -4 | 58 |
| Thalamus ventral lateral | R | 27 | 5.72 | 5.23 | .002 | 14 | -16 | 2 |
| Precentral gyrus | L | 38 | 5.59 | 5.13 | .004 | -50 | -6 | 50 |
| Rolandic Operculum | L | 20 | 5.54 | 5.09 | .005 | -62 | 4 | 8 |
| Caudate | R | 17 | 5.47 | 5.03 | .006 | 12 | 10 | 14 |
| Inferior frontal gyrus, triangular part | L | 29 | 5.18 | 4.81 | .016 | -40 | 32 | 4 |
| *PLA > NAL* |  |  |  |  |  |  |  |  |
| **Gyrus rectus/OFC medial** | **R** | **293** | **7.96** | **6.83** | **0.000** | **8** | **46** | **-20** |
| Gyrus rectus/OFC medial | R |  | 5.79 | 5.29 | 0.002 | 10 | 28 | -18 |
| **Superior frontal gyrus medial** | **L** | **50** | **6.73** | **5.99** | **0.000** | **-6** | **52** | **50** |
| **Superior frontal gyrus dorsolateral** | **L** | **21** | **5.72** | **5.23** | **0.002** | **-16** | **22** | **66** |
| **Superior frontal gyrus medial** | **L** | **56** | **5.66** | **5.19** | **0.003** | **-6** | **32** | **40** |
| **Middle frontal gyrus** | **L** | **95** | **5.48** | **5.05** | **0.006** | **-32** | **34** | **50** |
| Middle frontal gyrus | L |  | 5.38 | 4.96 | 0.008 | -30 | 32 | 42 |
| Superior frontal gyrus dorsolateral | L |  | 5.03 | 4.68 | 0.027 | -24 | 42 | 50 |
| **Middle frontal gyrus** | **L** | **23** | **5.34** | **4.93** | **0.009** | **-28** | **52** | **24** |
| **OFC lateral** | **R** | **16** | **5.32** | **4.92** | **0.010** | **52** | **40** | **-14** |
| **Superior frontal gyrus dorsolateral** | **L** | **21** | **5.15** | **4.78** | **0.018** | **-24** | **48** | **24** |
| Superior frontal gyrus dorsolateral | L |  | 5.12 | 4.75 | 0.020 | -24 | 56 | 16 |
| *PLA (food > touch) > AMI (food > touch)* |  |  |  |  |  |  |  |  |
| No suprathreshold clusters |  |  |  |  |  |  |  |  |
| *PLA (food > touch) > NAL (food > touch)* |  |  |  |  |  |  |  |  |
| No suprathreshold clusters |  |  |  |  |  |  |  |  |

Note. Hemi, hemisphere; k, cluster size (extended threshold k = 15); x y z, MNI coordinates; PLA, placebo; AMI, amisulpride; NAL, naltrexone. Brain regions were labelled with the Automated Anatomical Labelling Atlas (AAL) 3 (Rolls, Huang, et al., 2020).

